# *N*^6^ -methyladenosine modification and the YTHDF2 reader protein play cell type specific roles in lytic viral gene expression during Kaposi’s sarcoma-associated herpesvirus infection

**DOI:** 10.1101/201475

**Authors:** Charles Hesser, John Karijolich, Dan Dominissini, Chuan He, Britt Glaunsinger

## Abstract

Methylation at the *N*^6^ position of adenosine (m^6^A) is a highly prevalent and reversible modification within eukaryotic mRNAs that has been linked to many stages of RNA processing and fate. Recent studies suggest that m^6^A deposition and proteins involved in the m^6^A pathway play a diverse set of roles in either restricting or modulating the lifecycles of select viruses. Here, we report that m^6^A levels are significantly increased in cells infected with the oncogenic human DNA virus Kaposi’s sarcoma-associated herpesvirus (KSHV). Transcriptome-wide m^6^A-sequencing of the KSHV-positive renal carcinoma cell line iSLK.219 during lytic reactivation revealed the presence of m^6^A across multiple kinetic classes of viral transcripts, and a concomitant decrease in m^6^A levels across much of the host transcriptome. However, we found that depletion of the m^6^A machinery had differential pro- and anti-viral impacts on viral gene expression depending on the cell-type analyzed. In iSLK.219 and iSLK.BAC16 cells the pathway functioned in a pro-viral manner, as depletion of the m^6^A writer METTL3 and the reader YTHDF2 significantly impaired virion production. In iSLK.219 cells the defect was linked to their roles in the post-transcriptional accumulation of the major viral lytic transactivator ORF50, which is m^6^A modified. In contrast, although the ORF50 mRNA was also m^6^A modified in KSHV infected B cells, ORF50 protein expression was instead increased upon depletion of METTL3, or, to a lesser extent, YTHDF2. These results highlight that the m^6^A pathway is centrally involved in regulating KSHV gene expression, and underscore how the outcome of this dynamically regulated modification can vary significantly between cell types.

**Author Summary:** In addition to its roles in regulating cellular RNA fate, methylation at the *N*^6^ position of adenosine (m^6^A) of mRNA has recently emerged as a mechanism for regulating viral infection. While it has been known for over 40 years that the mRNA of nuclear replicating DNA viruses contain m^6^A, only recently have studies began to examine the distribution of this modification across viral transcripts, as well as characterize its functional impact upon viral lifecycles. Here, we apply m^6^A-sequencing to map the location of m^6^A modifications throughout the transcriptome of the oncogenic human DNA virus Kaposi’s sarcoma-associated herpesvirus (KSHV). We show that the m^6^A machinery functions in a cell type specific manner to either promote or inhibit KSHV gene expression. Thus, the KSHV lifecycle is impacted by the m^6^A pathway, but the functional outcome may depend on cell lineage specific differences in m^6^A-based regulation.

## Introduction

The addition of chemical modifications is critical to many steps of mRNA processing and the regulation of mRNA fate. There are more than 100 different RNA modifications, but the most abundant internal modification of eukaryotic mRNAs is *N*^6^-methyladenosine (m^6^A), which impacts nearly every stage of the posttranscriptional mRNA lifecycle from splicing through translation and decay [1-6]. The breadth of impacts ascribed to the m^6^A mark can be attributed to its creation of new platforms for protein recognition, in part via local changes to the RNA structure [4,7-12]. The reversibility of m^6^A deposition through the activity of demethylases termed erasers adds a further layer of complexity by enabling dynamic regulation, for example during developmental transitions and stress [1,4,5,13-15]. Deposition of m^6^A occurs co- or post-transcriptionally through a complex of proteins with methyltransferase activity known as writers, which include the catalytic subunit METTL3 and cofactors such as METTL14 and WTAP [1,4,14,16,17]. The modification is then functionally ‘interpreted’ through the selective binding of m^6^A reader proteins, whose interactions with the mRNA promote distinct fates.

The best-characterized m^6^A readers are the YTH domain proteins. The nuclear YTHDC1 reader promotes exon inclusion [6], whereupon m^6^A-containing mRNA fate is guided in the cytoplasm by the YTHDF1-3 readers. Generally speaking, YTHDF1 directs mRNAs with 3’ UTR m^6^A modifications to promote translation [3], whereas YTHDF2 recruits the CCR4-NOT deadenylase complex to promote mRNA decay [18]. YTHDF3 has been proposed to serve as a co-factor to potentiate the effects of YTHDF1 and 2 [3,19,20]. Although the individual effects of YTHDF1 and 2 seem opposing, the YTHDF proteins may coordinate to promote accelerated mRNA processing during developmental transitions and cellular stress [1]. YTHDC2, the fifth member of the YTH family proteins, was recently shown to play critical roles in mammalian spermatogenesis through regulating translation efficiency of target transcripts [21]. Additional examples of distinct functions for m^6^A readers under specific contexts such as heat shock are rapidly emerging [13].

Given the prevalence of the m^6^A modification on cellular mRNAs, it is not surprising that a number of viruses have been shown to contain m^6^A in their RNA [22-29]. Indeed, a potential viral benefit could be a less robust innate antiviral immune response, as m^6^A modification of in vitro synthesized RNAs diminishes recognition by immune sensors such as TLR3 and RIG-I [30,31]. That said, the functional consequences of viral mRNA modification appear diverse and include both pro- and anti-viral roles. In the case of Influenza A, a negative sense ssRNA virus, m^6^A and the reader YTHDF2 have been shown to promote viral replication [32]. Furthermore, multiple studies have mapped the sites of m^6^A modification in the human immunodeficiency virus (HIV) genome, and shown that it promotes the nuclear export of HIV mRNA as well as viral protein synthesis and RNA replication [24,26,28]. Roles for the YTHDF proteins in during HIV infection remain varied however, as Tirumuru and colleagues propose they function in an antiviral context by binding viral RNA and inhibiting reverse transcription, while Kennedy and colleagues observe they enhance HIV replication and viral titers [24,28]. A more consistently anti-viral role for the m^6^A pathway has been described for the *Flaviviridae*, whose (+) RNA genomes are replicated exclusively in the cytoplasm and contain multiple m^6^A sites in their genomic RNA [23,25]. An elegant study by Horner and colleagues showed that depletion of m^6^A writers and readers or the introduction of m^6^A-abrogating mutations in the viral E1 gene all selectively inhibit hepatitis C virus (HCV) assembly [23]. Similarly, depletion of METTL3 or METTL14 enhances Zika virion production [25].

Despite the fact that m^6^A modification of DNA viruses was first reported more than 40 years ago for simian virus 40, herpes simplex virus type 1, and adenovirus type 2, roles for the modification in these and other DNA viruses remain largely unexplored [33-37]. Unlike most RNA viruses, with few exceptions DNA viruses replicate in the nucleus and rely on the cellular transcription and RNA processing machinery, indicating their gene expression strategies are likely interwoven with the m^6^A pathway. Indeed, it was recently shown that the nuclear reader YTHDC1 potentiates viral mRNA splicing during lytic infection with Kaposi’s sarcoma-associated herpesvirus (KSHV) [38]. Furthermore, new evidence suggests m^6^A modification potentiates the translation of late SV40 mRNAs [39], further indicating that this pathway is likely to exert a wide range of effects on viral lifecycles.

Here, we sought to address roles for the m^6^A pathway during lytic KSHV infection by measuring and mapping the abundance of m^6^A marks across the viral and host transcriptome. This gammaherpesvirus remains the leading etiologic agent of cancer in AIDS patients, in addition to causing the lymphoproliferative disorders multicentric Castleman’s disease and primary effusion lymphoma. The default state for KSHV in cultured cells is latency, although in select cell types the virus can be reactivated to engage in lytic replication, which involves a temporally ordered cascade of gene expression. We reveal that m^6^A levels are significantly increased upon KSHV reactivation, which is due to a combination of m^6^A deposition across multiple kinetic classes of viral transcripts and a concomitant decrease in m^6^A levels across much of the host transcriptome. Depletion of m^6^A writer and cytoplasmic reader proteins impaired viral lytic cycle progression in the KSHV iSLK.219 and iSLK.BAC16 reactivation models, suggesting this pathway potentiates the KSHV lytic cycle. Interestingly, however, the roles for the m^6^A writer and readers shifted to instead display neutral or anti-viral activity in the TREX-BCBL-1 reactivation model. These findings thus demonstrate that while KSHV mRNAs are marked by m^6^A, the functional consequences of this mark can vary significantly depending on cell context, reinforcing both the functional complexity and dynamic influence of m^6^A.

## Results

### KSHV mRNA contains m^6^A modifications

Epitranscriptome mapping has revealed significant roles for the m^6^A pathway in the lifecycle and regulation of several RNA viruses, but at the time we initiated these studies, similar global analyses had yet to be performed for a DNA virus. Given that herpesviral mRNAs are transcribed and processed in the nucleus using the cellular RNA biogenesis machinery, we hypothesized that these viruses would engage the m^6^A pathway. We therefore first quantified how KSHV reactivation impacted total cellular m^6^A levels in the KSHV-positive renal carcinoma cell line iSLK.219 (**Fig 1**). These cells are a widely used model for studying viral lytic events, as they stably express the KSHV genome in a tightly latent state but harbor a doxycycline (dox)-inducible version of the major viral lytic transactivator ORF50 (also known as RTA) that enables efficient entry into the lytic cycle [40,41]. Polyadenylated (polyA+) RNA was enriched from untreated (latent) or dox-reactivated iSLK.219 cells and the levels of m^6^A were quantitatively analyzed by liquid chromatography-tandem mass spectrometry (LC-MS/MS) (**Fig 1A**). Indeed, we observed a three-fold increase in total m^6^A levels upon KSHV lytic reactivation, suggesting that m^6^A deposition significantly increased during viral replication (**Fig 1B**).

**Fig 1.**
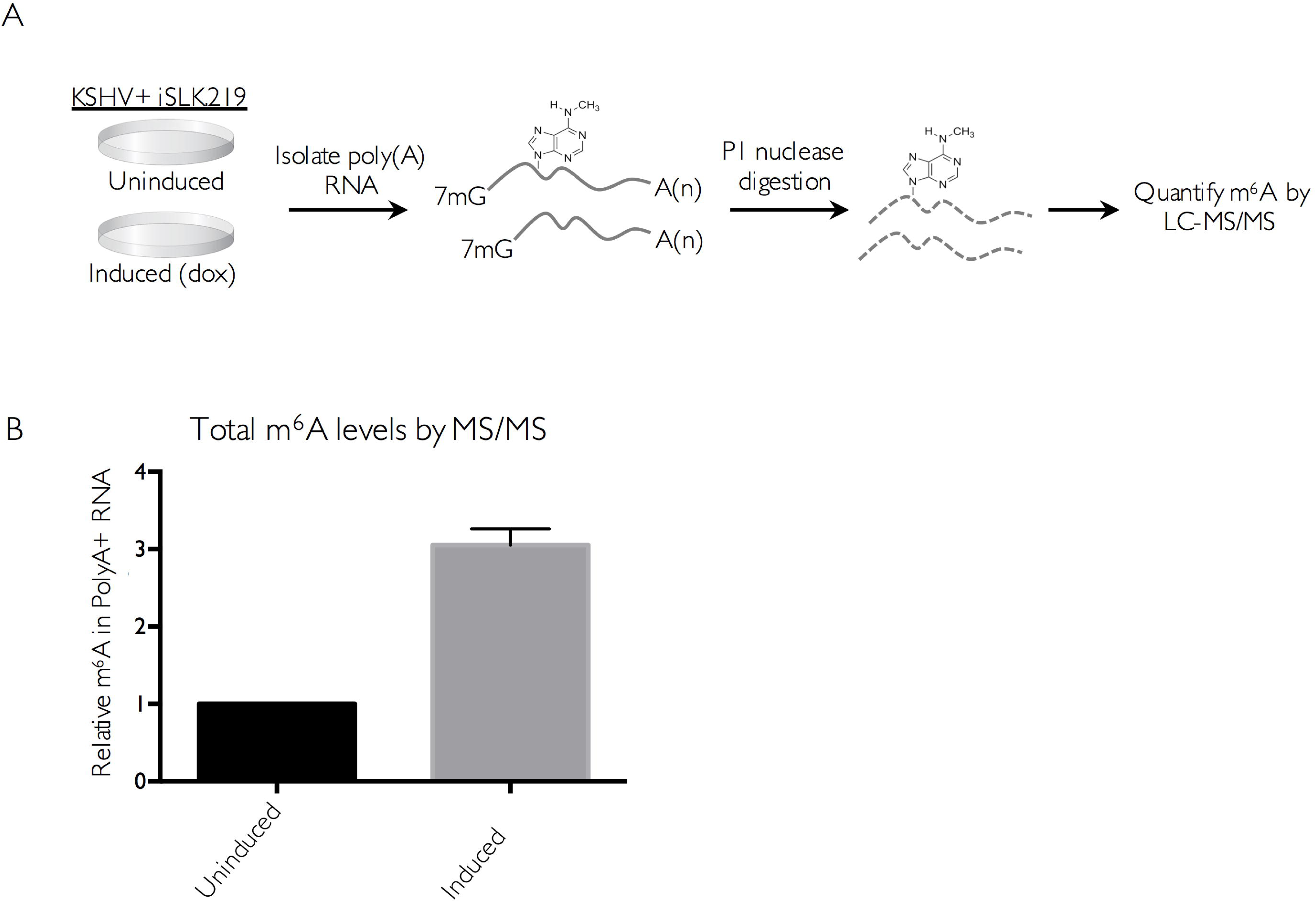
m^6^A increases upon KSHV reactivation. (A) Schematic of the experimental setup. iSLK.219 cells were induced with doxycycline for 5 days to induce the lytic cycle, and total RNA was collected and subjected to oligo dT selection to purify poly(A) RNA. Polyadenylated RNA was spiked with 10uM of 5-fluorouridine and digested with nuclease P1 and alkaline phosphatase, and subjected to LC-MS/MS analysis. (B) Relative m^6^A content in iSLK.219 cells. The induced sample was normalized with respect to the uninduced sample (set to 1).

We next sought to discern whether the increase in m^6^A during the KSHV lytic cycle favors host or viral mRNAs using high throughput m^6^A RNA sequencing (m^6^A-seq) [42]. This technique can reveal both the relative abundance and general location of m^6^A in KSHV and cellular mRNA. Total m^6^A containing RNA was immunoprecipitated from 2 biological replicates of latent or lytically reactivated iSLK.219 cells using an m^6^A-specific antibody. DNase-treated total mRNA was fragmented to lengths of 100 nt prior to immunoprecipitation and then subjected to m^6^A-seq. Total RNA-seq was run in parallel for each sample, allowing the degree of m^6^A modification to be normalized with respect to transcript abundance because the levels of many transcripts change upon viral lytic reactivation. Peaks with a fold-change four or higher (FC>4) and a false discovery rate of 5% or lower (FDR>5%) in both replicates were considered significant, although it is possible that additional transcripts detectably modified to lower levels or in a more dynamic manner may also be functionally regulated by m^6^A (complete list of viral peaks with FC>2 in **S1 Table**). In lytically reactivated samples, 10 transcripts comprising genes of immediate early, early, and late kinetic classes displayed significant m^6^A modification in both replicates (**Fig 2A** and **S1 Fig**). Within these KSHV mRNAs, m^6^A peaks were detected primarily in coding regions, although in some cases the location of a peak in a coding region overlaps with a UTR (**S1 Fig**). Furthermore, all but one peak contains at least one instance of the GG(m^6^A)C consensus sequence. While many of the modified viral transcripts contained only one m^6^A peak, multiple peaks were found in certain transcripts, including the major lytic transactivator ORF50 (**Fig 2B**). Of note, exon2 of ORF50 contained one m^6^A peak of FC>4 in replicate one, and three m^6^A peaks in replicate two, each of which have at least one m^6^A consensus motif, further increasing confidence that these peaks accurately represent m^6^A modified sites. Furthermore, the viral ncRNA PAN, which has been reported to comprise over 80% of nuclear PolyA+ RNA during lytic reactivation [43], contains FC>4 peaks in both replicates. Modification of PAN likely accounts for the marked three-fold increase in cellular m^6^A content observed upon lytic reactivation (**Fig 1B**). As anticipated given the restricted viral gene expression profile during latency, unreactivated samples had many fewer m^6^A containing viral mRNAs, with the only FC>4 peaks occurring in both replicates located in ORF4. Although ORF4 is a lytic transcript, its coding region overlaps with the 3’ UTR of K1, which is expressed during both the latent and lytic phases of the viral lifecycle (**Fig 2A, S1 Fig**) [44,45].

**Fig 2.**
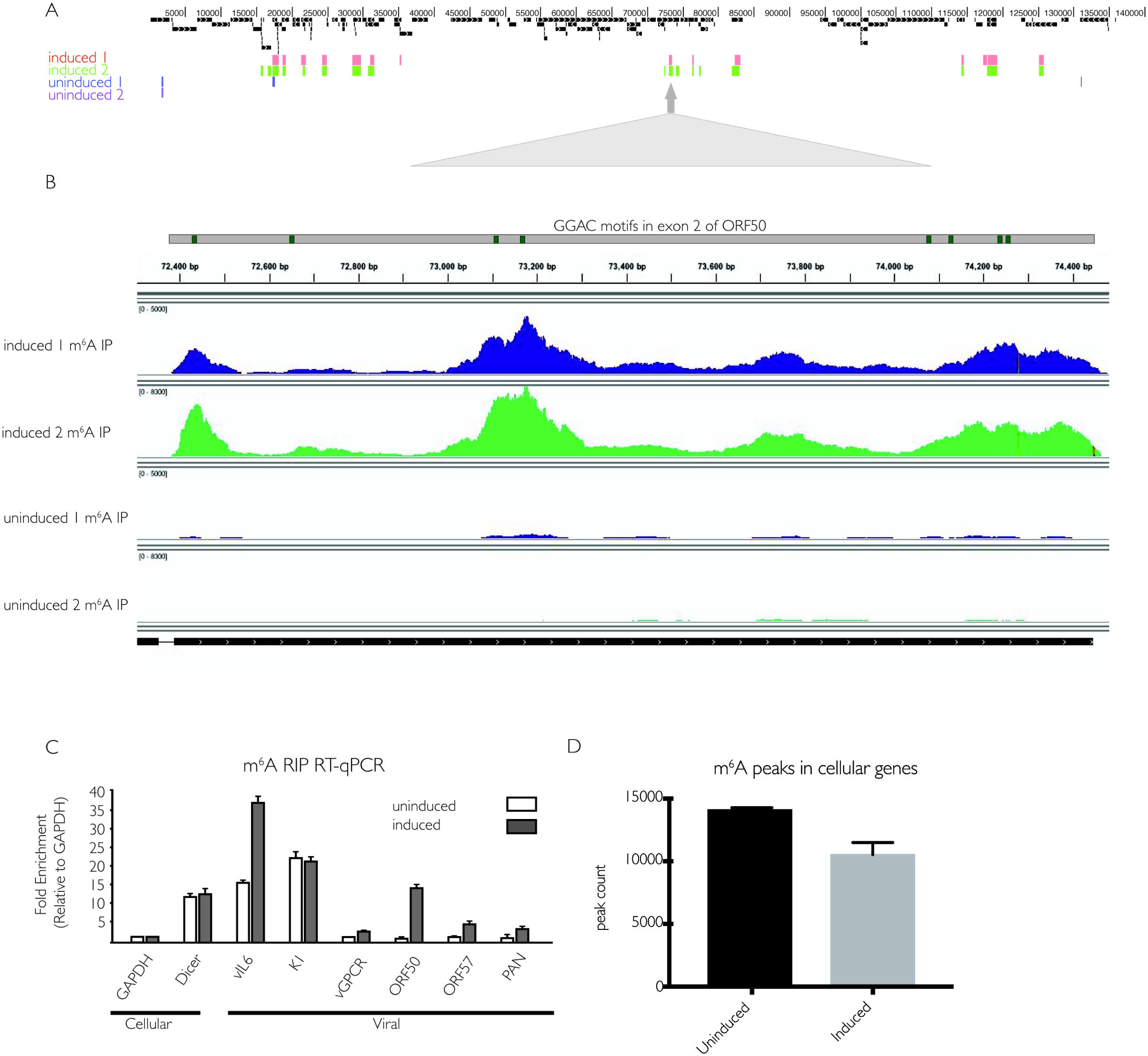
KSHV mRNA contains m^6^A modifications. (A) Two independent replicates of iSLK.219 cells containing latent KSHV were treated with dox for 5 days to induce the viral lytic cycle (induced) or left untreated to preserve viral latency (uninduced). DNase-treated RNA was isolated and subjected to m^6^Aseq. Displayed are peaks with a fold change of four or higher, comparing reads in the m^6^A-IP to the corresponding input. (B) Overview of sequencing reads from induced and uninduced m^6^A IP samples, aligned to the ORF50 transcript and the annotated GG(m^6^A)C consensus motifs found in exon 2 of ORF50. (C) Cells were induced as in (A), and total RNA was subjected to m^6^A RIP, followed by RT-qPCR using primers for the indicated viral and cellular genes. Values are displayed as fold change over input, normalized to GAPDH. (D) Quantification of cellular m^6^A peaks from m^6^Aseq analysis.

To validate the m^6^A-seq results, we performed m^6^A RNA immunoprecipitation (RIP) followed by quantitative real-time PCR (RT-qPCR) on six of the viral transcripts predicted to be m^6^A modified from the m^6^A-seq data. This technique allows determination of the relative level of m^6^A content in a given transcript compared to an unmodified transcript. As controls, we included primers for the cellular GAPDH transcript, which is known not to be m^6^A modified, and the Dicer transcript, which is m^6^A modified [42]. The m^6^A RIP RT-qPCR confirmed modification of the vIL-6, K1, ORF50, ORF57 and PAN viral transcripts, in agreement with m^6^A-seq results (**Fig 2C**). In summary, we found m^6^A modification in approximately one third of KSHV transcripts upon lytic reactivation, consistent with the hypothesis that this pathway contributes to KSHV gene expression.

We next compared the distribution of m^6^A peaks in host mRNAs from unreactivated versus reactivated cells to assess whether lytic KSHV infection altered the m^6^A profile of cellular transcripts. Analyzing the two independent replicates for each condition, we found an average of 14,092 m^6^A modification sites (FC>4 and FDR>5%) in host transcripts pre-reactivation, compared to 10,537 peaks post-reactivation (**Fig 2D** and **S2 Table**). We observed that this >25% decrease in m^6^A deposition on cellular mRNA encompassed a wide spectrum of transcripts, and no notable patterns were apparent by GO term analysis for functional categories enriched in the altered population. Thus, while the functional impact of the altered host m^6^A profile remains unresolved, the observation that KSHV lytic infection increased the level of m^6^A in total poly A+ RNA despite decreasing its presence in cellular mRNA implies that m^6^A deposition during infection favors viral transcripts.

### m^6^A and the reader YTHDF2 mediate viral gene expression and virion production in iSLK.219 cells

Given the significant deposition of m^6^A across KSHV transcripts, we reasoned that m^6^A might play an important role in potentiating the viral lifecycle. We therefore examined the effect of depleting the m^6^A writers and readers on KSHV virion production using a supernatant transfer assay. The KSHV genome in iSLK.219 cells contains a constitutively expressed version of GFP, which allows for fluorescence-based monitoring of infection by progeny virions. We performed siRNA-mediated knockdown of METTL3, the catalytic subunit responsible for m^6^A deposition, as well as the m^6^A readers YTHDF 1, 2 and 3 (**Fig 3A**). Cells were then treated with dox and sodium butyrate to induce lytic reactivation for 72 h, whereupon supernatants were collected and used to infect 293T recipient cells. The number of GFP positive 293T cells at 24 hpi was measured by flow cytometry (**Fig 3B**). Notably, for virus generated from METTL3 depleted cells, only 7% of recipient cells were infected compared to 82% for virus generated during treatment with a control siRNA (**Fig 3B**). YTHDF2 depletion caused an even more pronounced defect, resulting in a near absence of virion production (**Fig 3B**). In contrast, YTHDF3 knockdown resulted in only modest changes in virion production, while virion production was unaffected by YTHDF1 knockdown (**Fig 3B**). The prominent defect in virion production in METTL3 and YTHDF2 depleted cells was not due to knockdown-associated toxicity, as we did not observe changes in cell viability in siRNA treated cells (representative experiment shown in **S2 Fig**). Furthermore, we validated the results for YTHDF2 and YTHDF3 using independent siRNAs (**S2 Fig**). Thus, the m^6^A writer METTL3 and the reader YTHDF2 play important roles in driving KSHV infectious virion production in iSLK.219 cells.

**Fig 3.**
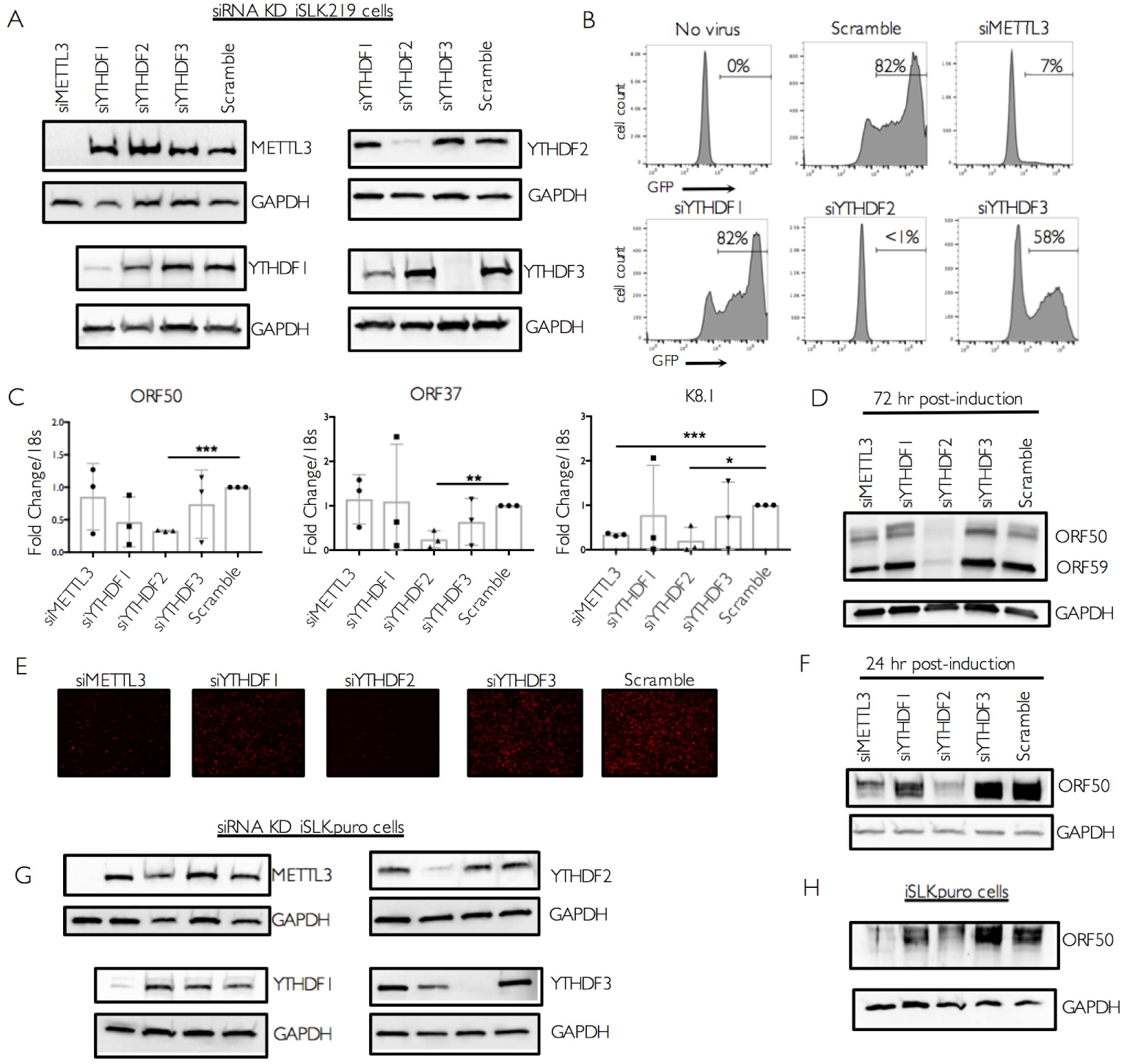
m^6^A and the reader YTHDF2 potentiate viral gene expression and virion production in iSLK.219 cells. Cells were transfected with control scramble (scr) siRNAs or siRNAs against METTL3, YTHDF1, 2, or 3, then reactivated for 72 hr with doxycycline and sodium butyrate. (A) Knockdown efficiency was measured by western blot using antibodies for the indicated protein, with GAPDH serving as a loading control in this and all subsequent figures. (B) Viral supernatant was collected from the reactivated iSLK.219 cells and transferred to uninfected HEK293T recipient cells. 24 h later, the recipient cells were analyzed by flow cytometry for the presence of GFP, indicating transfer of infectious virions. (C) ORF50, ORF37 and K8.1 gene expression was analyzed by RT-qPCR from cells treated with the indicated siRNAs. Data are from 3 independent experiments. Unpaired Student’s t test was used to evaluate the statistical difference between samples. Significance is shown for P values <0.05 (*), ≤ 0.01 (**), and ≤ 0.001 (***). (D) Expression of the viral ORF50 and ORF59 proteins in cells treated with the indicated siRNAs was measured by western blot 72hr post-reactivation. (E) Unreactivated iSLK.219 cells containing latent virus were treated with control scramble (scr) siRNAs or siRNAs targeting METTL3, YTHDF1, YTHDF2, or YTHDF3. The cells were then reactivated with dox and sodium butyrate for 48 hr and lytic reactivation was monitored by expression of the lytic promoter-driven red fluorescent protein. (F) Protein was harvested from the above described cells and subjected to western blot for ORF50 and the control GAPDH protein at 24 hr post-reactivation. (G-H) Uninfected iSLK.puro cells expressing DOX-inducible RTA were transfected with the indicated siRNAs for 48 h, then treated with dox for 24 hr to induce ORF50 expression. Knockdown efficiency (*G*) and ORF50 protein levels (*H*) were measured by western blot using antibodies for the indicated protein, with GAPDH serving as a loading control.

We then sought to determine the stage of the viral lifecycle impacted by the m^6^A pathway by measuring the impact of writer and reader depletion on the abundance of viral mRNAs of different kinetic classes. First, levels of representative immediate early, delayed early, and late viral mRNAs were measured by RT-qPCR following lytic reactivation for 72 hr. ORF50 and K8.1 transcripts contained at least one m^6^A peak, while ORF37 did not appear to be significantly modified in our m^6^A-seq data (see **S1 Table**). METTL3 depletion did not appear to impact accumulation of the ORF50 immediate early or ORF37 delayed early mRNAs at this time point, but resulted in a significant defect in accumulation of the K8.1 late gene mRNA (**Fig 3C**). Consistent with the virion production data, we observed a striking and consistent defect in the accumulation of each of the viral transcripts upon YTHDF2 depletion, suggesting that this protein is essential for lytic KSHV gene expression beginning at the immediate early stage (**Fig 3C**). Similar results were observed using an independent YTHDF2-targeting siRNA (**S2 Fig**). We also observed a prominent defect in accumulation of ORF50 and the delayed early ORF59 proteins by Western blot specifically upon YTHDF2 depletion (**Fig 3D**). In contrast, depletion of YTHDF1 or YTHDF3 did not reproducibly impact ORF50, ORF37, or K8.1 gene expression at 72 h post reactivation (**Fig 3C**).

In agreement with the above findings, we also observed that iSLK.219 cells depleted of METTL3 and YTHDF2 displayed a prominent defect in viral reactivation, as measured by expression of red fluorescent protein (RFP) driven by the PAN lytic cycle promoter from the viral genome (**Fig 3E**). Similarly, ORF50 protein production was also markedly reduced upon METTL3 or YTHDF2 depletion at the 24 h time point, which represents the early phase of the lytic cycle (**Fig 3F**).

To determine whether the effects of the m^6^A pathway on ORF50 were dependent on KSHV infection, we measured ORF50 protein in an uninfected iSLK cell line containing only the integrated, dox-inducible ORF50 gene (iSLK.puro cells) (**Fig 3G**). Similar to our findings with infected iSLK.219 cells, depletion of METTL3 or YTHDF2 strongly reduced ORF50 protein levels (**Fig 3H**). YTHDF3 depletion resulted in an increase in ORF50 expression, which we also observed to a more modest degree in the iSLK.219 cells (see **Fig 3D**). Collectively, these results suggest that m^6^A modification is integral to the KSHV lifecycle, and that YTHDF2 plays a particularly prominent role in mediating KSHV lytic gene expression in iSLK.219 cells. They further indicate that m^6^A modification can impact ORF50 expression in both uninfected and KSHV infected iSLK cells.

### The m^6^A pathway post-transcriptionally controls ORF50 expression in iSLK.219 cells, leading to a subsequent defect in transcriptional feedback at the ORF50 promoter

ORF50 is the major viral transcriptional transactivator, and its expression is essential to drive the KSHV lytic gene expression cascade [46]. The observations that ORF50 is m^6^A modified and that its accumulation is dependent on YTHDF2 indicate that the m^6^A pathway plays key roles in ORF50 mRNA biogenesis or fate in iSLK.219 cells, potentially explaining the lytic cycle progression defect in the knockdown cells. Deposition of m^6^A has been reported to occur both co-transcriptionally and post-transcriptionally [1,16,17,47]. To determine whether the m^6^A pathway is important for ORF50 synthesis or its posttranscriptional fate, we measured ORF50 transcription in reactivated iSLK.219 cells upon depletion of METTL3, YTHDF2, or YTHDF3 using 4-thiouridine (4sU) metabolic pulse labeling. 4sU is a uridine derivative that is incorporated into RNA during its transcription, and thiol-specific biotinylation of the 4sU-containing RNA enables its purification over streptavidin-coated beads [48,49]. At 24 h post reactivation, RNA in the siRNA treated iSLK.219 cells was pulse labeled with 4sU for 30 min, whereupon the labeled RNA was isolated by biotin-streptavidin purification and viral transcripts were quantified by RT-qPCR (**Fig 4A**). Despite the defect in ORF50 accumulation observed upon YTHDF2 depletion (see **Fig 3F**), we observed no decrease in 4sU-labeled ORF50 mRNA upon depletion of any of the m^6^A writer or reader proteins (**Fig 4B**). However, in YTHDF2 depleted cells, there was a prominent defect in the level of 4sU-labeled ORF37, likely because its transcription is dependent on the presence of ORF50 protein (**Fig 4C**).

**Fig 4.**
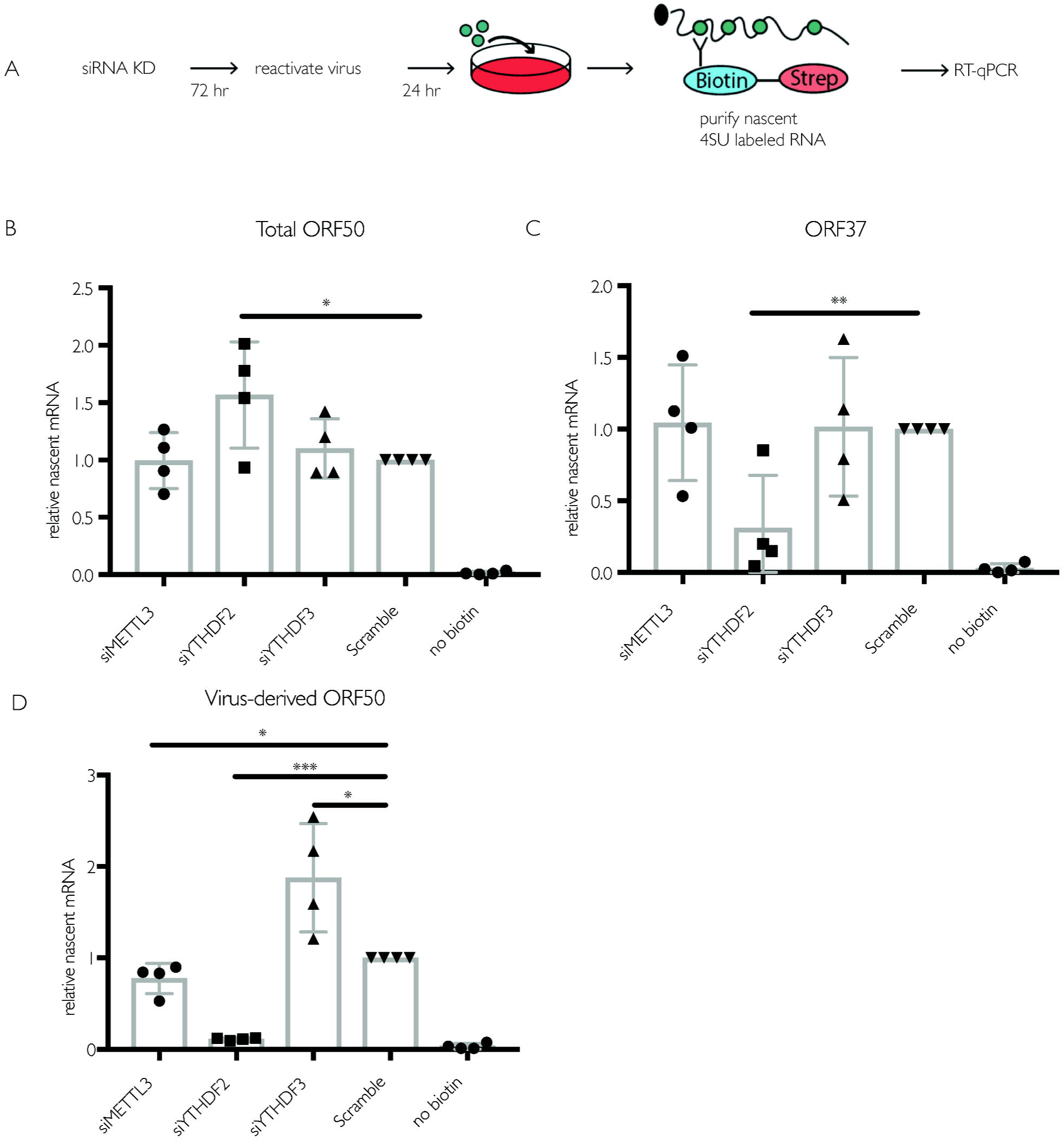
Depletion of the m^6^A writer and readers does not impact ORF50 nascent transcription in iSLK.219 cells. (A) Schematic of the experimental setup for measuring nascent RNA synthesis. Cells were transfected with the indicated siRNAs for 48h then reactivated for 24 hr with dox. 4sU was added for 30 minutes, whereupon 4sU-labeled RNA was isolated using biotin/streptavidin affinity purification, reverse transcribed, and analyzed by RT-qPCR using primers specific to ORF50 or ORF37. (B-D) Levels of 4sU-labeled total ORF50 (B), ORF37 (C), and ORF50 transcribed from the viral genome (virus-derived) (D) determined as described above. Unpaired Student’s t test was used to evaluate the statistical difference between samples. Significance is shown for P values <0.05 (*), ≤ 0.01 (**), and ≤ 0.001 (***).

The ORF50 mRNA detected in the above experiments represents a combination of the mRNA transcribed from the dox-inducible cassette as well as from the KSHV genome [41]. While the dox-inducible promoter is constitutively active under dox treatment, ORF50 transcription from KSHV is sensitive to ORF50 protein levels because it transactivates its own promoter [50]. The decreased ORF50 protein levels observed in Fig 3 might therefore lead to a selective reduction in transcription from the native ORF50 promoter by interfering with this positive transcriptional feedback. Indeed, primers designed to specifically recognize ORF50 derived from the viral genome revealed a marked defect in transcription of KSHV-derived ORF50 upon YTHDF2 depletion, as well as a slight reduction upon METTL3 depletion (**Fig 4D**). Collectively, the above results suggest that m^6^A initially functions to post-transcriptionally regulate ORF50 mRNA abundance, but that when ORF50 protein levels fall upon YTHDF2 or METTL3 knockdown, the positive transcriptional feedback mechanism at the viral promoter also becomes restricted.

### The impact of m^6^A on KSHV infection is cell type specific

To independently validate the METTL3 and YTHDF2 phenotypes, we also evaluated their importance in the iSLK.BAC16 model. Although independently generated, this is the same cell background as iSLK.219, including the dox-inducible ORF50, but instead contains the viral genome in the context of a bacterial artificial chromosome (BAC16) [51]. Similar to our results with the infected iSLK.219 cells, depletion of METTL3 or YTHDF2 in iSLK.BAC16 cells led to a significant defect in virion production as measured by supernatant transfer assays (**Fig 5A-C**). In addition, the total levels of ORF50 mRNA (from the dox-induced plus viral promoters) were unchanged between the different siRNA treated cells, while depletion of YTHDF2 led to a significant reduction in the level of BAC16-derived ORF50 and K8.1 mRNAs (**Fig 5D**). In contrast, METTL3 depletion did not significantly impact the level of ORF50, ORF37, or K8.1 transcripts. It should be noted that levels of METTL3 knockdown in excess of 80% have only been reported to reduce m^6^A levels in Poly A RNA by 20-30% [17]. Thus, at least some fraction of ORF50 (and other) transcripts may still be m^6^A modified due to residual enzyme activity of the remaining METTL3. In agreement with these observations, knockdown of METTL3 modestly reduced but did not eliminate the pool of m^6^A modified ORF50 or the cellular SON mRNAs in iSLK.BAC16 cells as measured by m^6^A RIP RT-qPCR (**S3 Fig**). Finally, we observed that although ORF59 protein levels were consistently reduced upon YTHDF2 knockdown, and to a more variable extent upon METTL3 depletion, we did not detect the same marked effects on ORF50 protein levels in iSLK.BAC16 cells as in iSLK.219 cells (**Fig 5E**). In summary, although iSLK.219 and iSLK.BAC16 cells exhibit a somewhat different gene expression profile in the context of YTHDF2 and METTL3 knockdown, depletion of these m^6^A pathway components restricts the KSHV lytic lifecycle in both models.

**Fig 5.**
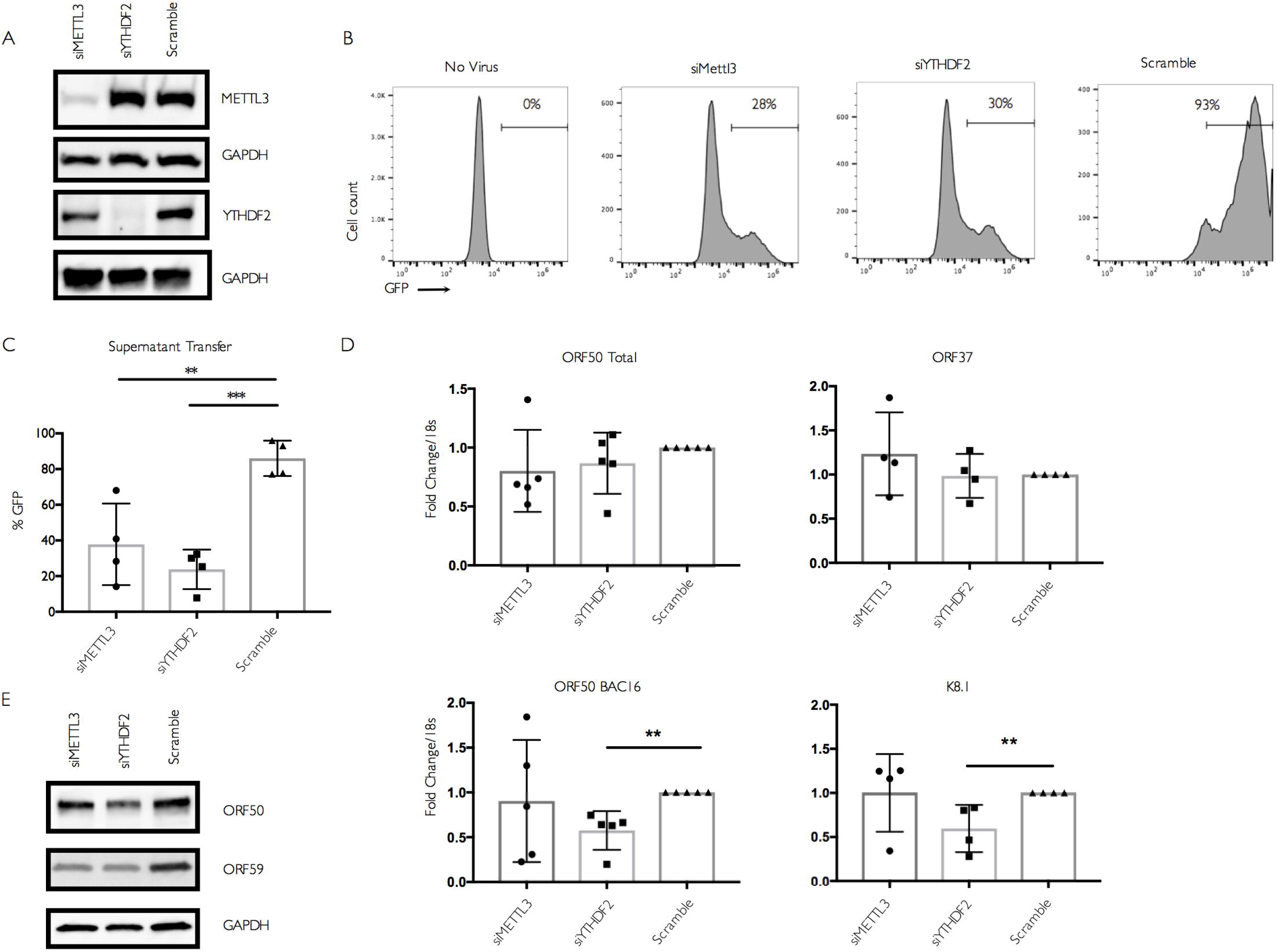
METTL3 and YTHDF2 are important for KSHV lytic replication in iSLK.BAC16 cells. Cells were transfected with control scramble (scr) siRNAs or siRNAs against METTL3 or YTHDF2, then reactivated for 24 hr with doxycycline and sodium butyrate. (A) Knockdown efficiency was measured by western blot using antibodies for the indicated protein. (B) Viral supernatant was collected from the reactivated iSLK.BAC16 cells 72 hr post-reactivation and transferred to uninfected HEK293T recipient cells. 24 h later, the recipient cells were analyzed by flow cytometry for the presence of GFP, indicating transfer of infectious virions. (C) Quantification of supernatant transfer results from four independent experiments. (D) ORF50, ORF37, and K8.1 gene expression 24 hr post-reactivation was analyzed by RT-qPCR from cells treated with the indicated siRNAs. Data for ORF50 are from five independent experiments, while ORF37 and K8.1 data are from four independent experiments. Unpaired Student’s t test was used to evaluate the statistical difference between samples in panels C-D. Significance is shown for P values ≤ 0.01 (**) and ≤ 0.001 (***). (E) Western blots showing expression of the viral ORF50 and ORF59 proteins at 24 hr post reactivation of iSLK.BAC16 cells treated with the indicated siRNAs.

Given the diversity of functions reported form^6^A in controlling cellular processes and virus infections [1,22,27], we also sought to evaluate the role of this pathway in mediating ORF50 expression in another widely used KSHV infected cell line of distinct origin, the B cell line TREX-BCBL-1 [52]. Similar to iSLK.219 and iSLK.BAC16 cells, TREX-BCBL-1 cells also contain a dox-inducible copy of ORF50 to boost reactivation. First, we evaluated whether the ORF50 transcript was m^6^A modified in TREX-BCBL-1 cells by m^6^A RIP, followed by RT-qPCR using control or ORF50 specific primers. Indeed, there was a clear enrichment of ORF50 in the reactivated, m^6^A-containing RNA population (**Fig 6A**). As expected, we detected the m^6^A modified DICER transcript in both reactivated and unreactivated cells, whereas the unmodified GAPDH transcript was present in neither sample (**Fig 6A**).

**Fig 6.**
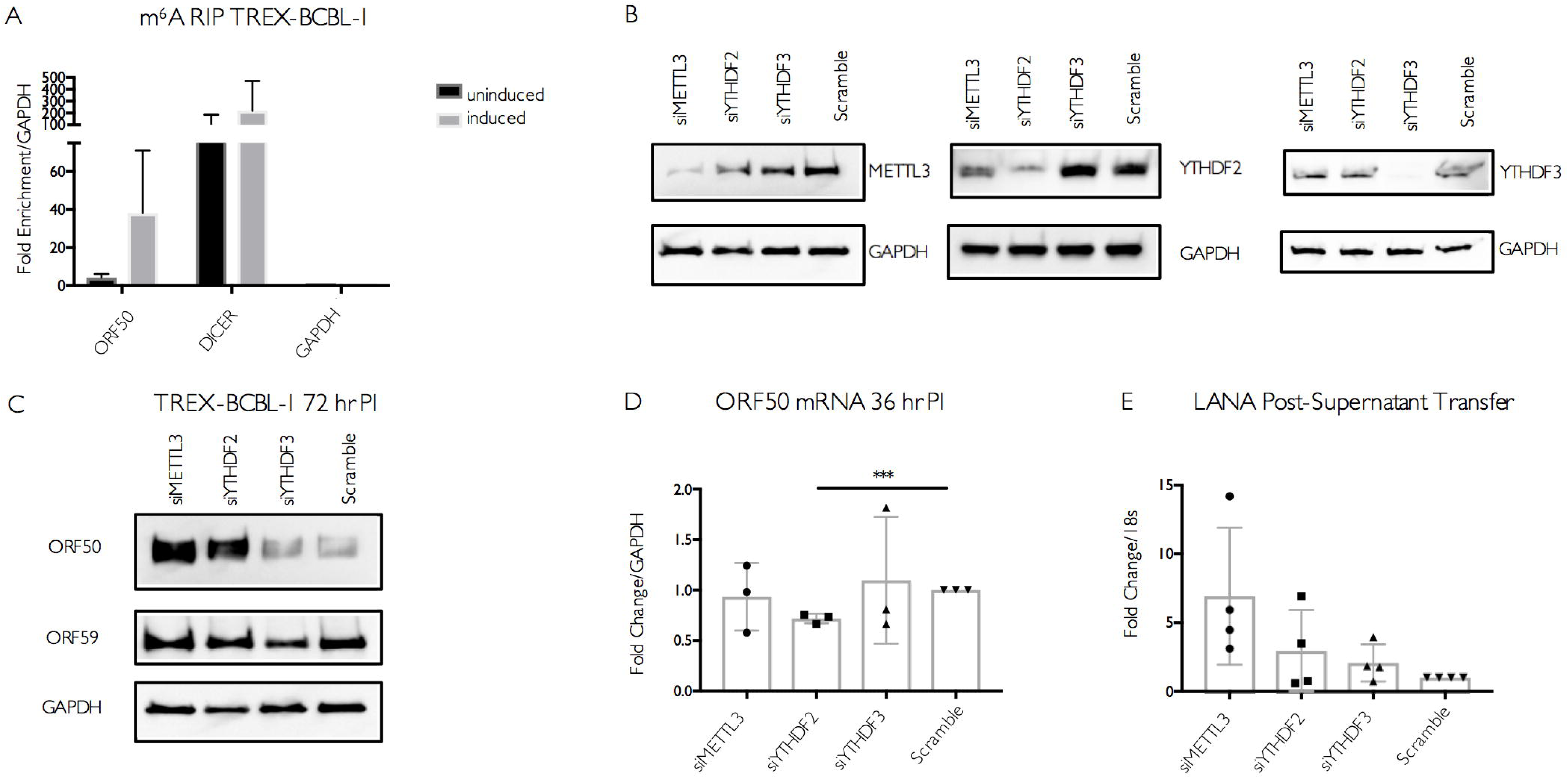
Increased viral gene expression upon m^6^A writer and reader depletion in TREX-BCBL-1 cells. (A) TREX-BCBL-1 cells were reactivated with dox for 72 hr, then total RNA was isolated and subjected to m^6^A RIP, followed by RT-qPCR for analysis of KSHV ORF50, GAPDH, and DICER. Values are displayed as fold change over input, normalized to the GAPDH negative control. Data are included from 3 biological replicates. (B-D) TREX-BCBL-1 cells were nucleofected with control scramble (scr) siRNAs or siRNAs specific to METTL3, YTHDF2, or YTHDF3, then lytically reactivated by treatment with dox, TPA and ionomycin for 72 hr. (B) Knockdown efficiency of the m^6^A proteins relative to the loading control GAPDH was visualized by western blot. (C) Levels of the KSHV ORF50 and ORF59 proteins were assayed by western blot in the control and m^6^A protein-depleted samples. Additional replicates are shown in **S4 Fig**. (D) ORF50 gene expression was analyzed by RT-qPCR from cells treated with the indicated siRNAs and reactivated for 36 hr with dox, TPA and ionomycin. (E) Viral supernatant from the reactivated control or m^6^A protein depleted TREX-BCBL-1 cells was transferred to uninfected HEK293T recipient cells, whereupon transfer of infection was quantified by RT-qPCR for the viral LANA transcript 48 hr post supernatant transfer. Individual data points represent 3 independent experiments. Unpaired Student’s t test was used to evaluate the statistical difference between samples. Significance is shown for P values <0.05, with *** representing P value ≤ 0.001.

The m^6^A pathway components were then depleted from TREX-BCBL-1 cells via siRNA treatment, whereupon cells were reactivated for 72 hr with dox, TPA, and ionomycin. As knockdown efficiency for YTHDF1 was inconsistent in this cell type, we focused on the impact of METTL3, YTHDF2, and YTHDF3. We observed no significant changes in the level of ORF50 mRNA upon METTL3 or YTHDF3 depletion (**Fig 6B, D**). Although there was a consistent decrease in ORF50 mRNA in the YTHDF2 depleted cells, this may be due to the fact that YTHDF2 knockdown modestly decreased the viability of TREX-BCBL1 cells (**S4 Fig**). Surprisingly, however, METTL3 knockdown and, to a more variable extent YTHDF2 knockdown, resulted in increased ORF50 protein expression (**Fig 6C**, additional replicate experiments showing ORF50 levels in **S4 Fig**). YTHDF3 depletion did not significantly impact ORF50 or ORF59 protein (**Fig 6C**). Thus, unlike in iSLK cells, METTL3 and YTHDF2 appear to restrict ORF50 expression in TREX-BCBL1 cells. These phenotypic differences were not due to distinct virus-induced alterations in the abundance of METTL3, YTHDF2, or YTHDF3, as levels of these proteins remained consistent following lytic reactivation in TREX-BCBL1, iSLK.219, and iSLK.BAC16 cells (**S5 Fig**).

Finally, to determine whether the m^6^A pathway components impacted the outcome of the viral lifecycle in TREX-BCBL1 cells, we measured the impact of METTL3, YTHDF2, and YTHDF3 protein knockdown on virion production using a supernatant transfer assay. TREX-BCBL-1 cells lack the viral GFP marker, and thus infection of recipient cells was instead measured by RT-qPCR for the KSHV latency-associated LANA transcript. Again in contrast to the iSLK cell data, we observed that METTL3, YTHDF2, and YTHDF3 were dispensable for virion production in TREX-BCBL-1 cells (**Fig 6E**). Instead, METTL3 depletion consistently resulted in a modest, though not statistically significant, increase in the level of LANA mRNA in the recipient cells (**Fig 6E**). In summary, METTL3 and YTHDF2 appear to function in a pro-viral capacity and promote ORF50 expression in iSLK.219 and iSLK.BAC16 cells, but instead restrict ORF50 expression in TREX-BCBL-1 cells. These findings highlight how at least a subset of m^6^A pathway functions and targets may diverge between cell types.

## Discussion

Although m^6^A modification of viral RNAs has been recognized for more than 40 years, only recently are the contributions of this epitranscriptomic mark towards viral life cycles beginning to be revealed. Thus far, global epitranscriptomic analyses have documented m^6^A deposition during infections with KSHV, SV40, HIV, Influenza A virus and several members of the *Flaviviridae*, with a diverse set of resulting pro- and anti-viral roles [23-26,28,32,39,53]. The breadth and occasionally apparently contrasting functions for the m^6^A pathway during infection are perhaps unsurprising given the dynamic role for this modification in controlling mRNA fate and its ability to impact virtually every stage of host gene expression [1,27]. Our global analysis of the m^6^A epitranscriptome during lytic infection with the DNA virus KSHV showed the presence of m^6^A across multiple kinetic classes of viral transcripts and a general decrease in m^6^A deposition on cellular mRNAs. In the widely used KSHV-positive cell lines iSLK.219 and iSLK.BAC16, we found that depletion of several components of the m^6^A pathway inhibited the KSHV lytic cycle, most notably in iSLK.219 cells by restricting accumulation of the viral lytic transactivator ORF50. The YTHDF2 reader protein proved particularly important, as its depletion eliminated lytic entry and virion production. These observations are suggestive of a pro-viral role for m^6^A in the iSLK.219 and iSLK.BAC16 KSHV reactivation models. However, m^6^A marks on mRNA in a cell are widespread and contribute to a large variety of cellular and pathogenic processes that likely occur in a cell type or context-dependent manner. In this regard, it is notable that a distinct set of phenotypes was observed for m^6^A pathway components in the B cell line TREX-BCBL-1. Here, depletion of METTL3 and YTHDF2 increased ORF50 abundance, more suggestive of an anti-viral role. Thus, although KSHV engages the m^6^A pathway in multiple cell types, these findings underscore the importance of not broadly extrapolating m^6^A roles from a particular cell type, as this complex regulatory pathway can functionally vary in a cell type dependent manner.

What might be the basis for these phenotypic differences between cell types in the context of KSHV infection? m^6^A deposition was also recently reported in many KSHV mRNAs in BCBL-1 cells, including ORF50 [38]. Furthermore, while this work was in revision, Tan and colleagues documented extensive modification of KSHV transcripts during latent KSHV infection of multiple cell types, as well as upon lytic infection of iSLK.BAC16 and TREX-BCBL-1 cells [53]. Notably, while numerous differences were found in the cellular m^6^A profiles between the two cell lines, many peaks in viral transcripts were consistent across cell types, including two out of three m^6^A peaks in ORF50 [53]. In agreement with these studies, we also observed extensive modification of KSHV mRNAs, and observed that ORF50 is modified in iSLK.BAC16 cells, iSLK.219 cells and TREX-BCBL-1 cells. Thus, it is not the case that the viral mRNAs engage the m^6^A methyltransferase machinery in one cell type but not the other, although it is clear that site specificity of m^6^A deposition, particularly in host mRNAs, can vary between cell lines. The facts that m^6^A deposition is dynamic and does not strictly occur on consensus motifs render this possibility challenging to resolve. Indeed, how m^6^A deposition selectively controls gene regulation on particular transcripts or under particular stimuli remains a central unanswered question in the field [1]. We hypothesize that the distinct phenotypes derive either from how the viral modifications are ‘interpreted’ in each cell type and/or indirect effects driven by an altered m^6^A profile on cellular mRNAs. The recent finding that m^6^A modification of ORF50 in BCBL-1 cells contributes to efficient splicing through binding of YTHDC1 argues that modifications can have a direct cis-acting impact on KSHV mRNA fate [38]. However, herpesviral mRNAs are heavily reliant on host machinery at every stage of their biogenesis. Given that cellular mRNA fate is significantly altered upon depletion of METTL3 and the YTHDF reader proteins [1-3,18,54], it is possible that cell type specific changes in the abundance of a host factor(s) required for viral mRNA stability also contribute to the phenotypic differences. Furthermore, in HIV infected cells m^6^A modification and YTHDF proteins have been proposed to have a combination of pro-viral and anti-viral effects, including negatively impacting reverse transcription, enhancing mRNA export, and increasing viral protein production [24,26,28]. Therefore, the m^6^A pathway might similarly facilitate distinct phenotypes at different stages of the KSHV lifecycle.

Although our m^6^A-seq results are in agreement with the recent report from Tan and colleagues, our data on the role of YTHDF2 in iSLK.BAC16 cells differs from theirs [53]. They did not evaluate the impact of METTL3 depletion, but reported that YTHDF2 depletion increased KSHV replication in these cells. In contrast, we observed a significant reduction in virion production upon depletion of YTHDF2 in both iSLK.219 and iSLK.BAC16 cells. Given the similarity in approaches used to evaluate the impact of YTHDF2, the basis for these differences remains unclear. However, our experiments comparing the iSLK.219 and iSLK.BAC16 cells indicates that even in cell lines of the same origin there can be differences in the m^6^A-associated viral gene expression signatures.

As the ‘interpreters’ of m^6^A marks, the individual reader proteins play prominent roles in modulating gene expression. Generally speaking, in HeLa and 293T cells, YTHDF1 binding correlates with increased translational efficiency, YTHDF2 binding accelerates mRNA decay, and YTHDF3 may serve as a cofactor to assist the other reader protein function [1-4,19,20,54]. However, other roles for these factors are rapidly emerging, particularly in the context of cell stress, infection, or in the control of specific transcripts [7,13,15,23-26,28,54,55]. Furthermore, m^6^A is enriched in certain tissues, and different m^6^A patterns have been found depending on the tissue and developmental stage [42,56]. Intriguingly, a recent study showed that hypoxia increases global m^6^A content of mRNA, with many m^6^A modified RNAs exhibiting increased stability, raising the possibility that m^6^A deposition could also stabilize transcripts during other forms of cellular stress [57]. In KSHV-infected iSLK.219 cells, YTHDF2 appears essential for the post-transcriptional accumulation of ORF50, a role seemingly at odds with its more canonical mRNA destabilizing function. In this regard, it was recently revealed that SV40 late transcripts contain multiple m^6^A sites, and that YTHDF2 strongly promotes SV40 replication [39]. Thus, YTHDF2 has been shown to play a pro-viral role in the context of both DNA and RNA viruses. Although we observed less dramatic viral gene expression phenotypes upon METTL3 depletion, it nonetheless was required for WT levels of progeny virion production in iSLK.219 and iSLK.BAC16 cells. An important consideration may be that m^6^A factors differentially impact specific KSHV transcripts, or play different roles at distinct times during infection. However, dissecting these possibilities is likely to be complicated by the changes in ORF50 expression (either positive or negative), which will have ripple effects on the entire lytic life cycle. Another relevant question is the extent to which m^6^A mediates its effects on KSHV gene expression co-transcriptionally versus post-transcriptionally. A recent report indicated that m^6^A is primarily installed in nascent mRNA in exons and affects cytoplasmic stability, but not splicing [16,47]. It has also been demonstrated that that m^6^A can be installed co-transcriptionally, and that slowing the rate of RNA Pol II elongation enhances m^6^A modification of mRNAs in a manner that ultimately decreases translation efficiency [16,47]. These add to a growing body of literature indicating that the position of m^6^A in a transcript is a key feature impacting the functional consequence of the modification [1,3,6,7,13,20]. For example, m^6^A in the 3’ UTR has been shown to recruit YTHDF1 and enhance translation initiation in HeLa cells, while deposition of m^6^A in the 5’ UTR has been shown to enhance 5’ cap independent translation [3,7,13]. Whether these position-linked effects on translation extend to viral transcripts remains to be tested, although there does not appear to be a consistent enrichment in a particular region of viral mRNAs for the viruses analyzed thus far. In KSHV, m^6^A sites are found throughout viral ORFs, some of which also overlap with untranslated regions of other viral transcripts. As KSHV transcription depends on the host RNA Pol II, the speed of transcriptional elongation on viral mRNAs likely impacts co-transcriptional deposition and positioning of m^6^A, and thus may ultimately regulate translation efficiency of a given mRNA. Thus, in the context of KSHV reactivation, a wide variety of mechanisms exist through which m^6^A modification could impact the transcription and translation of viral mRNA. Deciphering these remains an important challenge for future studies, as we are currently in the early stages of understanding how this and other viruses interface with the m^6^A RNA modification pathway.

## Methods and Methods

### Cell culture

The renal carcinoma cell line iSLK.puro containing a doxycycline-inducible copy of ORF50, and the KSHV infected renal carcinoma cell lines iSLK.219 and iSLK.BAC16 bearing doxycycline-inducible ORF50 [41] were cultured in Dulbecco’s modified Eagle medium (DMEM; Invitrogen) with 10% fetal bovine serum (FBS; Invitrogen, HyClone) and 100 U/ml penicillin-streptomycin (Invitrogen). The KSHV-positive B cell line TREX-BCBL-1 containing a doxycycline-inducible version of ORF50 [52] was cultured in RPMI medium (Invitrogen) supplemented with 20% FBS, 100 U/ml penicillin/streptomycin, and 200 uM L-glutamine (Invitrogen). HEK293T cells (ATCC) were grown in DMEM (Invitrogen) supplemented with 10% FBS. To induce lytic reactivation of iSLK.219 cells, 2×10^6^ cells were plated in a 10 cm dish with 1 μg/ml doxycycline (BD Biosciences) and 1 mM sodium butyrate for 72 hr. Lytic reactivation of TREX-BCBL-1 cells was achieved by treatment of 7×10^5^ cells/ml with 20 ng/ml 2-O-tetradecanoylphorbol-13-acetate (TPA, Sigma), 1 μg/ml doxycycline (BD Biosciences), and 500 ng/ml ionomycin (Fisher Scientific) for 72 hr (western Blot blots for viral gene expression), or for 120 hr (supernatant transfer experiments).

### siRNA experiments

For iSLK.219 cells, 100 pmol of siRNA was reverse transfected into 5×10^5^ cells plated in a 6-well dish using Lipofectamine RNAimax (Life Technologies). 24 hr post transfection, cells were trypsinized and re-seeded on a 10 cm plate. The next day, a second transfection was performed on the expanded cells with the same concentration of siRNA (400 pmol siRNA and 2×10^6^ cells). The following day, cells were lytically reactivated in a 10 cm plate. 24 hr post-reactivation, cells were lysed in RIPA buffer (10 mM Tris-Cl (pH 8.0), 1 mM EDTA, 1% Triton X-100, 0.1% sodium deoxycholate, 0.1% SDS, 140 mM NaCl) to evaluate knockdown efficiency. siRNA experiments in iSLK.BAC16 cells were conducted with the same siRNAs and concentrations. For experiments to assess mRNA and protein levels at 24 hr post-reactivation, one round of siRNA knockdown was performed 48 hr prior to reactivation, and knockdown efficiency was evaluated at the time of cell harvest. For iSLK.BAC16 supernatant transfer experiments, two rounds of siRNA treatment were used, as described for iSLK.219 cells.

For TREX-BCBL-1 cells, 200 pmol of siRNA was nucleofected into 2×10^6^ cells using Lonza Cell Line Nucleofector Kit V and a Lonza Nucleofector 2b set to Program T001. After nucleofection, cells were immediately resuspended in 2.2 ml of RPMI media in a 12 well plate. 48 hr later, 200 pmol of siRNA was added again to 2×10^6^ cells using the same protocol. 48 hr after the second transfection, cells were lysed in RIPA buffer and knockdown efficiency was analyzed by Western Blot. Cell viability post-nucleofection was assessed using a Countess II Automated Cell Counter (Life Technologies) with Trypan blue staining. For RT-qPCR experiments, two rounds of siRNA knockdown were performed under the identical conditions, except using an Invitrogen Neon Nucleofector with a single pulse of 1350 volts and pulse length of 40 ms.

The following Qiagen siRNAs were used: SI00764715 and SI04279121 targeting YTHDF1, SI04205761 targeting YTHDF3, custom siRNA targeting METTL3 (sequence targeted: CTGCAAGTATGTTCACTATGA). The following Dharmacon siRNAs were used: SMARTpool siGENOME (M-021009-01-0005), targeting YTHDF2, and siGENOME Non-Targeting siRNA Pool #1 (D0012061305). These same siRNAs were used in all three cell lines for the experiments in Figs 3-6. In addition, independent siRNAs (Qiagen SI04174534 targeting YTHDF2, Qiagen SI00764778 targeting YTHDF3 and Qiagen SI03650318 (negative control siRNA)) were used in S2 Fig.

### Supernatant transfer assay and quantification of virion production

Assays were performed as previously described [58]. Briefly, for iSLK.219 and iSLK.BAC16 cells, viral supernatant was collected 72 hr post-reactivation, filtered, and added to uninfected HEK293T cells by spinfection (1500 rpm, 90 minutes at room temperature). 12 hr later, supernatant was removed and replaced with fresh media, whereupon the cells were assessed for the successful transfer of the GFP-containing KSHV BAC 24 hr post-infection using a BD Accuri C6 flow cytometer. Briefly, cells were trypsinized, fixed in 4% paraformaldehyde, washed twice in PBS and resuspended in FACS Buffer (PBS with 1% FBS). Uninfected HEK293T cells were used to define the GFP negative population. The percentage of GFP expressing cells was quantified using FlowJo Software (FlowJo LLC). For virus produced in TREX-BCBL-1 cells, supernatant transfers were performed as in iSLK.219 cells, except the virus was transferred to HEK293T cells at 120 hr post-reactivation. To quantify virus produced in TREX-BCBL-1 cells, RNA was extracted from HEK293T cells 48 hr post-supernatant transfer, and viral gene expression was quantified by RT-qPCR using primers specific for LANA.

### Affinity purification and Western blotting

Cell lysate was collected and analyzed as previously described [58]. Briefly, iSLK.219, iSLK.BAC16 or TREX-BCBL-1 cells were trypsinized, washed with PBS and lysed in RIPA buffer with protease inhibitors. After washing, 4X Laemmli sample buffer (Bio-Rad) was added to samples to elute bound proteins. Lysates were resolved by SDS-PAGE and western blots were carried out with the following antibodies: rabbit ORF50 (gift of Yoshihiro Izumiya, UC Davis), rabbit α-K8.1 (1:10000, antibody generated for this study), rabbit α-ORF59 (1:10000, antibody generated for this study), rabbit α-METTL3 (Bethyl, 1:1000), rabbit α-YTHDF1 (Proteintech, 1:1000), rabbit α-YTHDF2 (Millipore, 1:1000), rabbit α-YTHDF3 (Sigma, 1:1000), and goat α-mouse and goat α-rabbit HRP secondary antibodies (1:5000; Southern Biotech).

### 4sU Labeling

Following siRNA knockdown and 24 hr reactivation, iSLK.219 cells were pulse labeled with DMEM containing 500 μM 4sU (Sigma) for 30 minutes, followed by PBS wash and immediate isolation of total RNA with TRIzol. 4sU isolation was performed as previously described [59]. 4sU isolated RNA was analyzed by RT-qPCR.

### RT-qPCR

Total RNA was harvested using TRIzol following the manufacturer’s protocol. Samples were DNase treated using Turbo DNase (Ambion), and cDNA was synthesized from 2 μg of total RNA using AMV reverse transcriptase (Promega), and used directly for quantitative PCR (qPCR) analysis with the DyNAmo ColorFlash SYBR green qPCR kit (Thermo Scientific). All qPCR results were normalized to levels of 18s (except GAPDH where indicated) and WT or scramble control set to 1. RT-qPCR primers used in this study are listed in **S4 Table**.

### LC-MS/MS analysis of m^6^A

Total RNA was isolated from iSLK219 cells with TRIzol reagent. Dynabeads mRNA purification kit (Ambion) was used to isolate polyA(+) RNAs from 100 μg of total RNA according to the manufacturer. 100-200 ng of polyadenylated RNA was spiked with 10 μM of 5-fluorouridine (Sigma) and digested by nuclease P1 (1 U) in 25 μL of buffer containing 25 mM NaCl and 2.5 mM ZnCl_2_ at 42 °C for 2-4 h, followed by addition of NH_4_HCO_3_ (1 M, 3 μL) and bacterial alkaline phosphatase (1 U) and incubation at 37 °C for 2 h. The sample was then filtered (Amicon 3K cutoff spin column), and 5 μL of the flow through was analyzed by liquid chromatography (LC) coupled to an Orbitrap-XL mass spectrometer (MS) equipped with an electrospray ionization source (QB3 Chemistry facility).

### m^6^A-RIP and m^6^A-RIP-sequencing

Total cellular RNA (containing KSHV RNA) was extracted and purified by TRIzol and then DNAse treated with Turbo DNase (Ambion). 30 μl protein G magnetic beads (Invitrogen) were blocked in 1% BSA solution for 1 hour, followed by incubation with 12.5 μg affinity-purified anti-m^6^A polyclonal antibody (Millipore) at 4°C for 2 hr with head-over-tail rotation. 100 μg purified RNA was added to the antibody-bound beads in IP buffer (150 mM NaCl, 0.1% NP-40, and 10 mM Tris-HCl [pH 7.4]) containing RNAse inhibitor and protease inhibitor cocktail and incubated overnight at 4°C with head-over-tail rotation. The beads were washed three times in IP buffer, and then RNA was competitively eluted with 6.7 mM m^6^A-free nucleotide solution (Sigma Aldrich). RNA in the eluate was phenol chloroform extracted and then reverse transcribed to cDNA for Real-Time qPCR analysis.

High-throughput sequencing of the KSHV methylome (m^6^A-seq) was carried following the previously published protocol [60]. In brief, 2.5 mg total cellular RNA was prepared from iSLK.219 cells that were either unreactivated, or reactivated for five days with doxycycline. RNA was isolated and DNAse treated as in the m^6^A RIP, except the RNA was first fragmented to lengths of ∼100 nt prior to immunoprecipitation with anti-m^6^A antibody (Synaptic Systems). Immunoprecipitated RNA fragments and comparable amounts of input were subjected to first-strand cDNA synthesis using the NEBNext Ultra RNA Library Prep Kit for Illumina (New England Biolabs). Sequencing was carried out on Illumina HiSeq2500 according to the manufacturer’s instructions, using 10 pM template per sample for cluster generation, TruSeq SR Cluster kit v3 (Illumina), TruSeq SBS Kit v3-HS (Illumina) and TruSeq Multiplex Sequencing primer kit (Illumina). A reference human transcriptome was prepared based on the University of California, Santa Cruz (UCSC) and a reference KSHV transcriptome based on KSHV 2.0 annotation [61]. Analysis of m^6^A peaks was performed using the model-based analysis of ChIP-seq (MACS) peak-calling algorithm. Peaks were considered significant if their MACS-assigned fold change was greater than four and individual FDR value less than 5%. Sequencing data are available on GEO repository (accession number GSE104621). Raw reads and alignment to the viral genome are shown in **S3 Table**.

## Statistical Analysis

All results are expressed as means +/-S.E.M. of experiments independently repeated at least three times, except where indicated. Unpaired Student’s t test was used to evaluate the statistical difference between samples. Significance was evaluated with P values <0.05.

## Acknowledgements

This research was funded by NIH (http://www.nih.gov/) grants R01AI122528 to B.G. and HG008688 to C.H. B.G. and C.H. are investigators of the Howard Hughes Medical Institute. We wish to thank Divya Nandakumar for her assistance with the m^6^A-seq read mapping, as well as all members of the Glaunsinger and Coscoy labs for helpful comments and suggestions.

## Supporting Information

**S1 Fig. Location of union FC>4 peaks within KSHV transcriptome**. Overview of sequencing reads aligned to regions of the KSHV transcriptome containing m^6^A modifications. Depicted are peaks with a fold change of four or higher in both replicates, comparing reads in the m^6^A-IP to the corresponding input. The blue and purple bars denote the sequences encompassed by the FC>4 peaks in each uninduced replicate (A), while the red and green colored bars denote the FC>4 peaks in each induced replicate (B). In the reference transcriptome, grey bars indicate annotated 3’UTRs or ncRNAs, while burgundy arrows depict ORFs. Note that the alignment of the ORF50 induced peaks can be found in Figure 2.

**S2 Fig**. (A) Independent siRNAs against YTHDF2 and YTHDF3 yield similar results as those shown in Fig 3. Cells were transfected with control scramble (scr) siRNAs (Qiagen SI03650318) or siRNAs against YTHDF2 (Qiagen SI04174534) or YTHDF3 (Qiagen SI00764778), then reactivated for 72 hr with doxycycline and sodium butyrate. Viral supernatant was collected from the reactivated iSLK.219 cells and transferred to uninfected HEK293T recipient cells. 24 h later, the recipient cells were analyzed by flow cytometry for the presence of GFP, indicating transfer of infectious virions. Data are from 2 independent experiments, with each replicate shown. (B) ORF50 and ORF37 gene expression was analyzed by RT-qPCR from the above cells at the time of supernatant transfer. (B) Viability of iSLK.BAC16 and iSLK.219 cells following siRNA transfection. Cells were transfected with the indicated siRNAs for 48 h, followed by lytic reactivation with dox and sodium butyrate for 48 h. Cells were collected and diluted 1:1 with Trypan blue prior to counting on a Countess II Automated Cell Counter. One representative experiment is shown.

**S3 Fig. Impact of METTL3 depletion on isolation of** ^**6**^**A modified mRNA in iSLK**.**BAC16 cells**

iSLK.BAC16 cells were subject to siRNA knockdown using METTL3 or control siRNA for 48 hr. Cells were reactivated for 24 hr with dox. (A) Western blot for knockdown efficiency at time of harvest. (B) Total RNA from harvested cells was then subject to m^6^A RIP RT-qPCR for the viral transcript ORF50 and cellular transcripts SON (m^6^A modified) and GAPDH (unmodified). Data shown are from 5 independent experimental replicates.

**S4 Fig**. (A) Quantification of cell viability following siRNA nucleofection and reactivation in TREX-BCBL-1 cells. TREX-BCBL-1 cells were nucleofected twice with the indicated siRNAs as described in the methods, and then reactivated for 36 hr with dox, PMA and ionomycin. Cells were collected and diluted 1:1 with Trypan blue prior to counting on a Countess II Automated Cell Counter. Viability from three independent experiments is depicted in the bar graphs. Unpaired Student’s t test was used to evaluate the statistical difference between samples. Significance is shown for P values <0.05 (*). (B) Western blots from replicate experiments showing viral ORF50 and ORF59 protein levels in TREX-BCBL-1 cells treated with the indicated siRNAs and reactivated with dox, TPA, and ionomycin as described in Fig. 6C. (C) Western blots showing viral ORF50 and ORF59 protein levels in TREX-BCBL-1 cells treated with the indicated siRNAs for 72 h prior to reactivation with TPA and ionomycin.

**S5 Fig. No changes in the levels of writers and readers following KSHV lytic reactivation**

iSLK.BAC16, iSLK.219 or TREX-BCBL-1 cells were reactivated where indicated with dox for 24 or 48 hr, at which point cells were harvested and lysates were analyzed by Western blot for METTL3, YTHDF2, YTHDF3, and the GAPDH loading control.

S1. Table: Full list of FC>2 peaks within KSHV transcripts in induced and uninduced samples.

S2. Table: Full list of FC>4 peaks within host transcripts in induced and uninduced samples.

S3. Table: Read counts and alignment to the KSHV genome.

S4. Table: List of RT-qPCR primers used in this study.

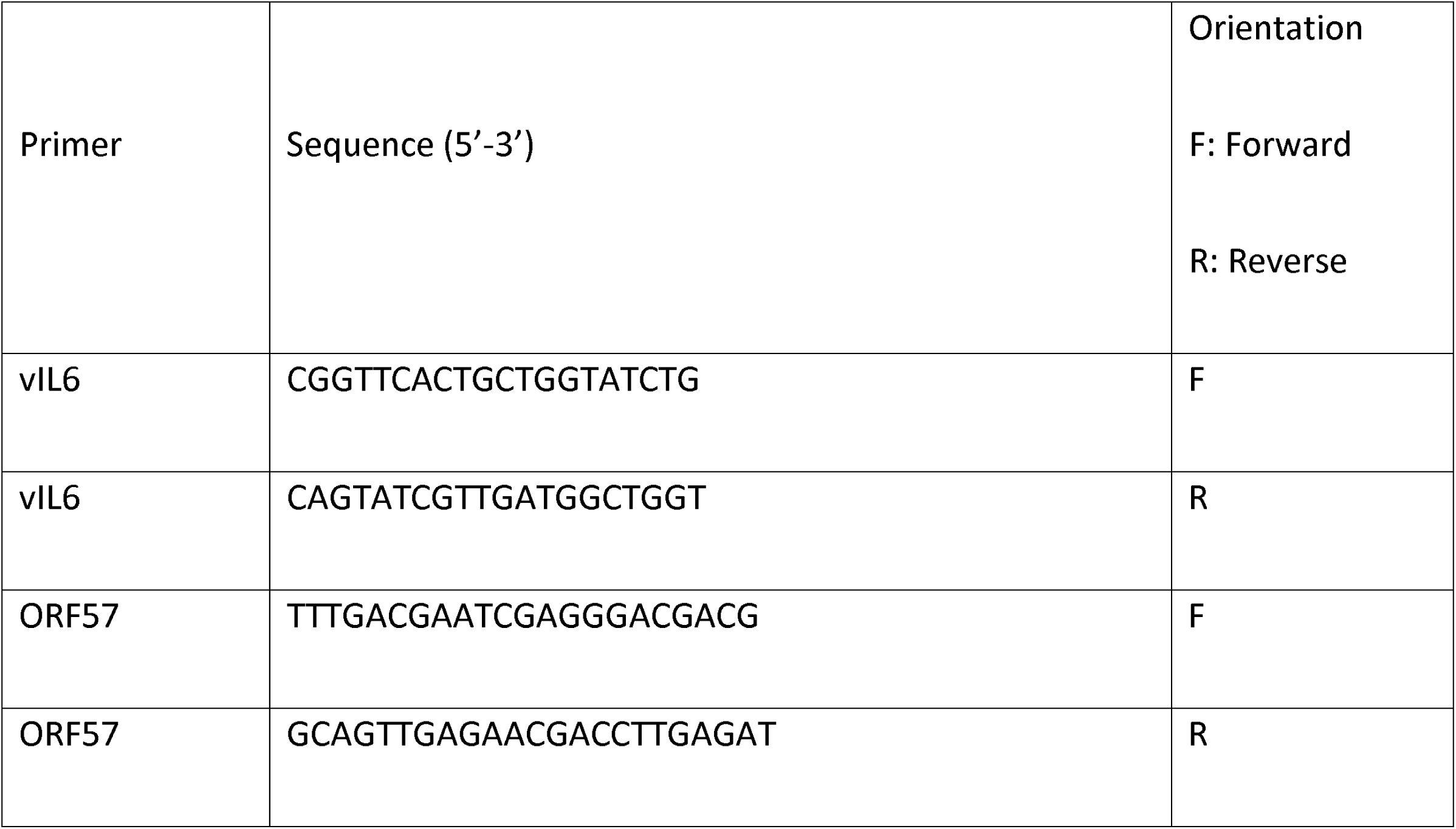

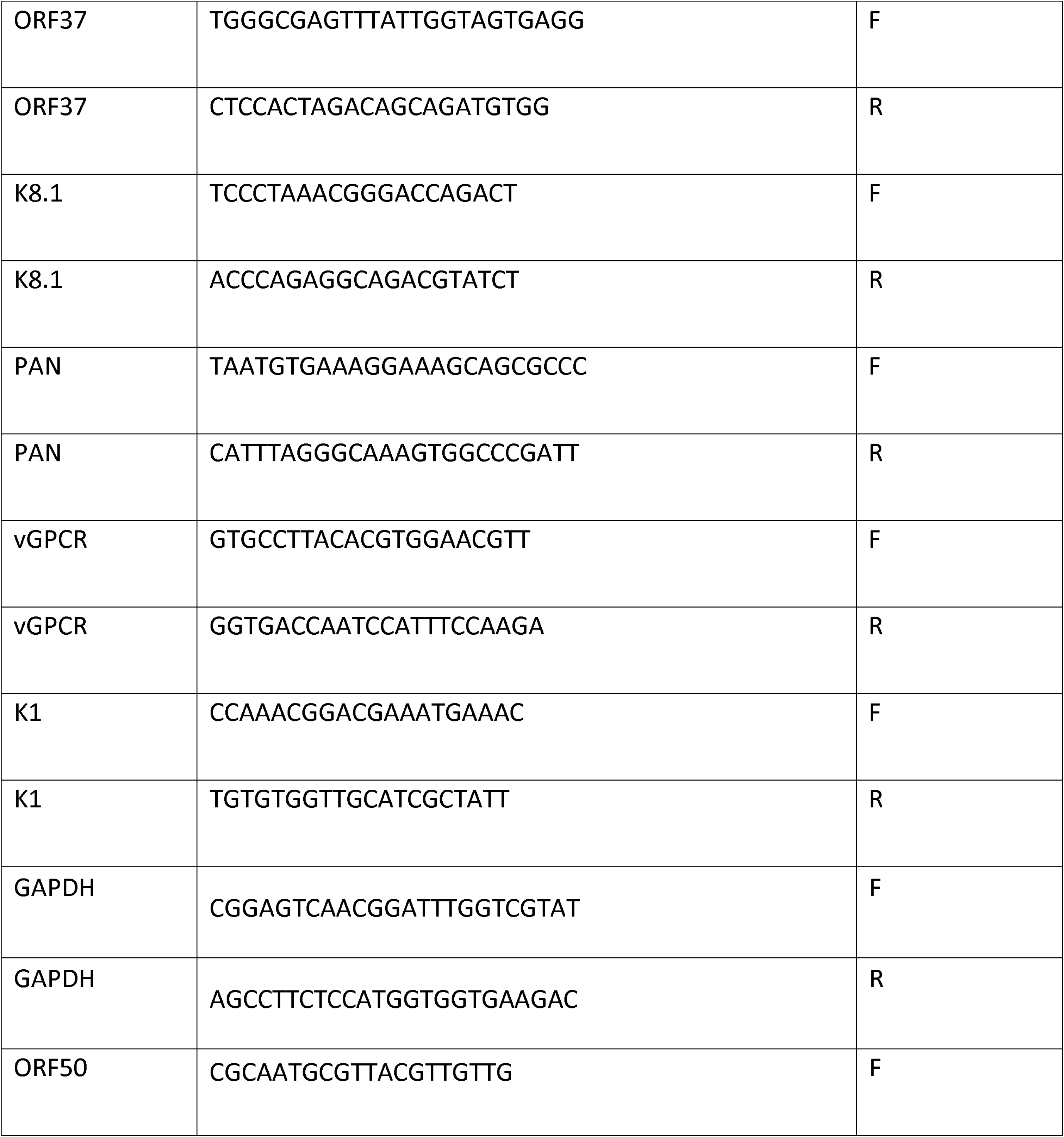

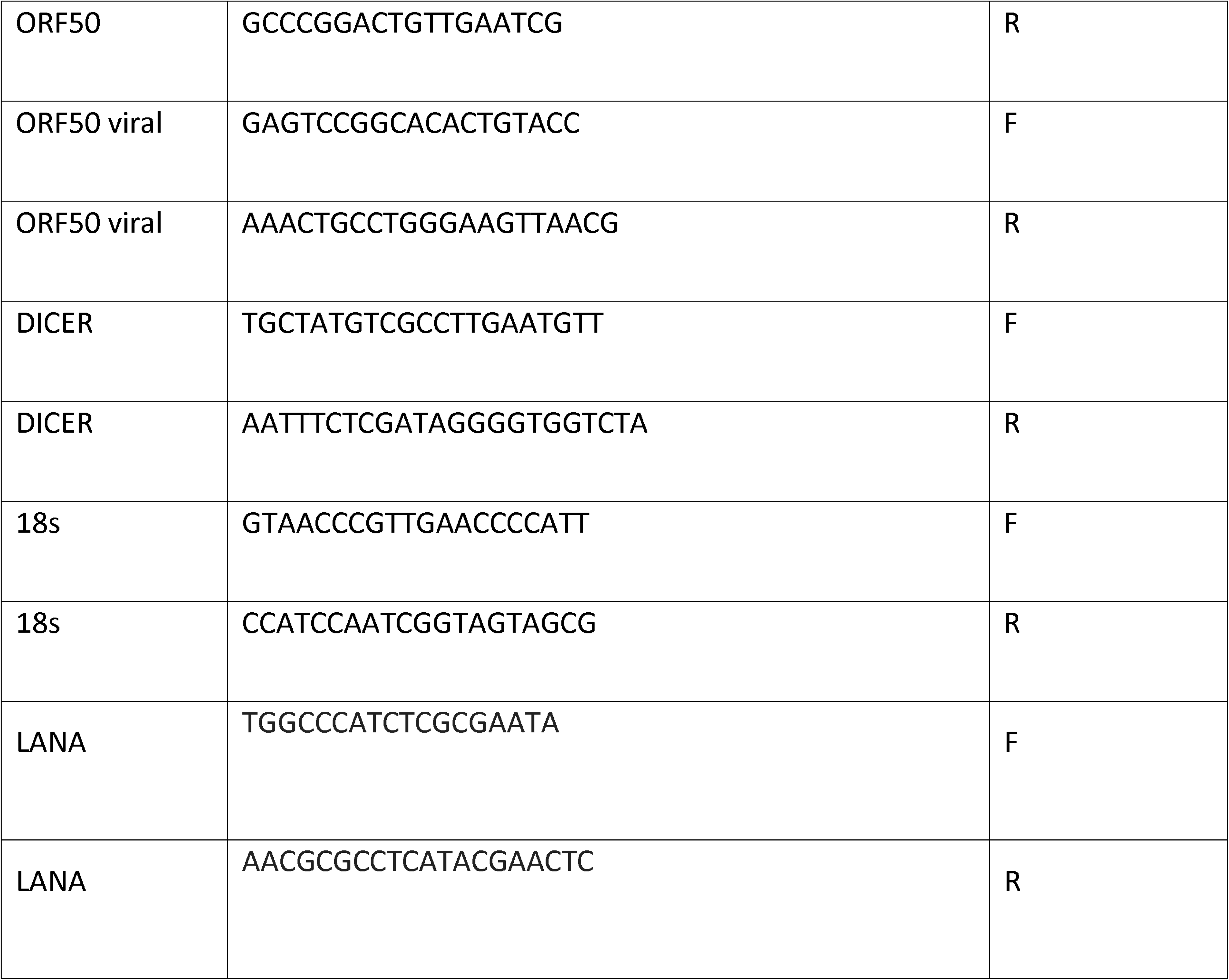

